# Variants of relative frequency methods for determining transcranial magnetic stimulation motor threshold do not provide accurate estimation of threshold

**DOI:** 10.64898/2026.07.27.741072

**Authors:** Boshuo Wang, Stefan M. Goetz, Angel V. Peterchev

## Abstract

Relative frequency methods such as five-out-of-ten have been used for a long time to determine motor threshold of transcranial magnetic stimulation (TMS), despite known severe limitations both in accuracy and speed. Variants of relative frequency methods have been developed to improve performance, such as adaptive staircasing and the recent RMT-Finder using binary search. However, they do not solve the fundamental problem underlying the use of relative frequencies to approximate the 50% TMS response rate at threshold. We demonstrate that the probability of false decisions using relative frequency due to at least half the pulses resulting in responses at subthreshold intensities or more than half the pulses resulting in no responses at suprathreshold intensity cannot be sufficiently reduced unless an impractically high pulse count is used, which substantially increases the duration of the thresholding procedure. Simulations in a virtual population of 25,000 subjects reveal that the accuracy of relative frequency methods is not improved by the adaptive staircasing or binary search variants. Although these variants reduce the number of pulses required to reach an estimate compared to the conventional relative frequency method, they are still slower and less accurate than more advanced thresholding methods such as stochastic approximation or maximum likelihood estimation. Therefore, these faster and more accurate methods should become the standard for motor thresholding of TMS, and the use of methods based on relative frequency should be discouraged.

**Highlights:**

- TMS motor thresholding should not use relative-frequency methods.
- Relative-frequency methods have a high probability of making false decisions.
- Variants of relative-frequency methods do not improve threshold accuracy.
- Stochastic approximation or maximum-likelihood estimation are faster and more accurate.

## 1. Introduction

The motor threshold (MT) of transcranial magnetic stimulation (TMS) is a critical parameter for neurological assessment of motor function and stimulation dosing of neuromodulation. Although many thresholding methods have been developed and evaluated over the years (Wang et al., 2023), the relative-frequency methods based on recommendations of the International Federation of Clinical Neurophysiology (IFCN) (Rossini et al., 1994, 2015) are still commonly used today due to their simplicity. The threshold is determined as the lowest stimulation intensity that generates response for at least half of the stimulation samples for a predetermined number of pulses, with the most commonly used method, five-out-of-ten, requiring between five and ten pulses at each intensity. Given the stochasticity of the motor responses, approximating the response probability by a relative frequency within just ten pulses can be rather imprecise (Awiszus, 2012, 2014), and even using a larger number of pulses per stimulation amplitude, e.g., twenty, may not substantially improve the accuracy of the threshold estimate (Wang et al., 2023).

Efforts to improve the relative-frequency methods include the adaptive staircasing variants (Stokes et al., 2005, 2013; Varnava et al., 2011) and the recent binary search variants used in “RMT-Finder” (Boidequin et al., 2026). Instead of adjusting the stimulation intensity with a small, fixed step size, typically 1% of maximum stimulator output (MSO), the staircasing variants titrate the intensity with decreasing step sizes in alternating directions based on the response counts. For example, one variant reduces the intensity by 5% MSO from an initial suprathreshold intensity until less than five responses are observed within ten pulses, then increases in steps of 2% MSO until five or more response are present, and finally decreases in steps of 1% MSO like the conventional IFCN relative-frequency methods (Stokes et al., 2013; Varnava et al., 2011). Another variant has an additional stage of stepping up by 10% MSO from subthreshold intensity at the beginning (Stokes et al., 2005). To improve on the staircasing methods, the binary search variants start with upper and lower bounds and iteratively reduce the range by testing the mid-point and moving either the lower or upper bound to the mid-point based on whether response count is less than five or not (Boidequin et al., 2026). The default variant (“auto”) uses 20% MSO and 90% MSO as the bounds and a “fast” variant incorporates participant-specific information by setting the bounds at 10% MSO below and above the individual’s hotspot search intensity. Here, we show that these variants suffer from the same issue of the basic relative frequency method and do not improve the accuracy of threshold estimates.

## 2. Probability of false decision

For any relative frequency method, a false decision is made when the intensity is subthreshold but at least half of the limited number of pulses result in responses and the intensity is incorrectly declared as suprathreshold, or vice versa when the intensity is suprathreshold but more than half of the applied pulses fail to elicit a response. A false decision results in early termination before the threshold is reached or late termination when the intensity steps below threshold for the IFCN method, reverses the stepping direction too early for staircasing method, and narrows the search space to the incorrect half not containing the threshold for binary search methods. Numerical analysis (details in supplement) shows that the false decision probability of the commonly used five-out-of-ten relative frequency is above 30% and up to 60% for intensities near the threshold (Fig. 1A, colored solid lines for *N* = 10). Although the probability drops for intensities further from the threshold, it remains at non-negligible values for intensities at the bounds of the ±5% relative error range, which defines clinical safety (Rossi et al., 2009; Awiszus, 2011). Especially, the probability of false suprathreshold decisions is inherently higher due to the protocol of relative frequency counting. For example, at an intensity of 95% MT, the false-decision probability is 6.4%, four times higher than the 1.5% false-decision probability at 105% MT. Although these values at the ±5% bounds may appear modest, the higher false-decision probability closer to threshold can easily step the intensity away from the threshold and out of the bounds. For a participant with a 50% MSO threshold, for example, the false-decision probabilities respectively are 37% for a false subthreshold decision at threshold or 33% and 12% for false suprathreshold decision at just 1% MSO and 2% MSO below threshold (98% and 96% MT). This is especially problematic for participants with low thresholds, as each 1% MSO step corresponds to a larger change relative to threshold. These false-decision probability near threshold can only be reduced to negligible levels by substantially increasing the pulse number at each intensity (Fig. 1A, dashed and dotted lines respectively for *N* = 20 and 40).

**Figure 1:**
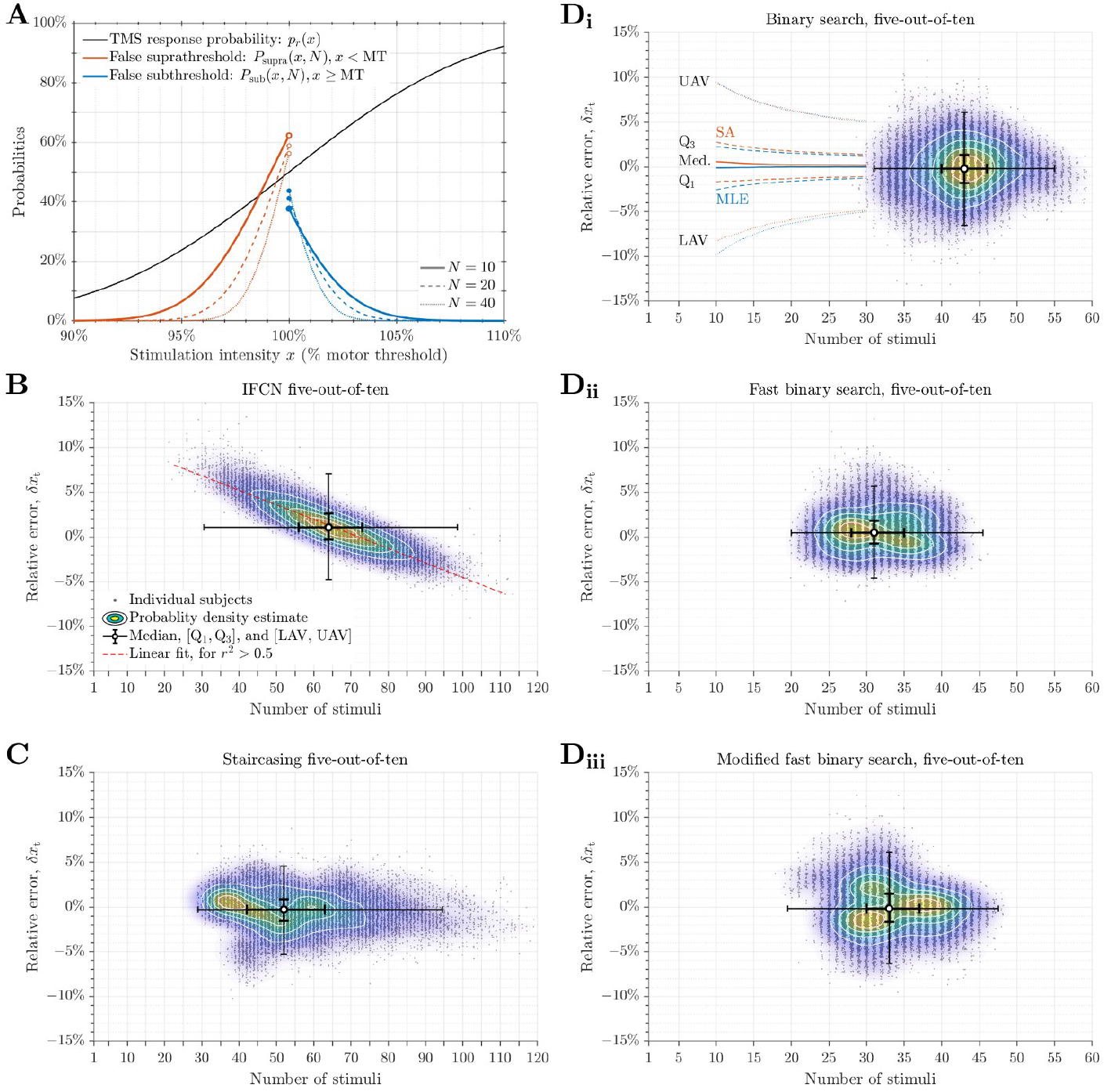
**A. Probability of false decisions for relative frequency methods as a function of stimulation intensity.** For typical TMS input–output response probabilities following a sigmoidal shape (black line), the probabilities of making false subthreshold decisions (blue lines, including left ends) and false suprathreshold decisions (orange lines, excluding right ends) are highest near threshold and decrease with distance from threshold. They only reduce to negligible levels outside the ±5% relative error range for the typical five-out-of-ten method (solid lines), especially for false suprathreshold decisions, or inside this range for higher pulse counts (dashed and dotted lines). **B–D. Relative threshold errors of the relative-frequency methods**. The relative errors *δx*_t_ are plotted against the number of stimuli required as dots for all the 25,000 virtual subjects. Several statistics are shown, with median as white circle, interquartile range between first and third quartiles (Q_1_ and Q_3_) as thick error bars, and ranges extending to the lower (LAV) and upper adjacent values (UAV)—which are 1.5 times the interquartile range below Q_1_ and above Q_3_—as thin error bars. A probability density estimate based on a normal kernel function is also shown with the colored contour map. Linear regression is shown for the IFCN relative-frequency methods as red lines. **B**. IFNC five-out-of-ten method, reproduced with data from Wang et al. (2023). **C**. Staircasing method. **D**. Binary search methods: **i**. default variant (“auto”), **ii**. fast variant, and **iii**. modified fast variant. Note different horizontal scales in panels B and C versus D_i_, D_ii_, and D_ii_. Panel D_i_ also shows the error statistics of maximum likelihood estimation (MLE, blue) and adaptive-stepping stochastic approximation (SA, orange) methods—which generate threshold estimates at each pulse—between pulse number 10 and 30, reproduced with data from Wang et al. (2023): median as thick lines, Q_1_ and Q_3_ as dashed lines, and LAV and UAV as dotted lines.

## 3. Bounds of binary search

Unlike the staircasing methods, the binary search relative frequency method assumes without testing that the initial bounds contain the threshold and there is no mechanism to extend the bounds if the intensity reaches one of them (Boidequin et al., 2026). For the “auto” variant, the wide range covered by the initial bounds based on prior data would likely be sufficient for most subjects, although the bounds may need to be adjusted for different stimulation pulse shape, TMS coils, and target muscles. Still, the fixed starting amplitude, 55% MSO, could be too high for participants with low thresholds (e.g., over 180% MT for a 30% MSO threshold). The subject-specific bounds for the “fast” variant, however, risk having the lower bound being above the threshold: as the hotspot search intensity should consistently elicit responses (Rossini et al., 2015), it could be more than 10% MSO higher than threshold, especially for target muscles with shallow response input−output curves (Koponen et al., 2024).

## 4. Performance of relative frequency variants

We used a realistic MEP model (Goetz et al., 2019) and a virtual population of 25,000 subjects (Wang et al., 2023) to simulate the performance of several variants of relative frequency method, including the staircasing method with the stepping sequence of −5%, +2%, and −1% MSO, and the binary search method with default bounds and its fast variant with participant-specific bounds (Boidequin et al., 2026). The hotspot intensity was defined as an intensity with a 95% response rate and, compared to the threshold, was on average 8.7% ± 0.7% MSO (range: 6.4% to 16.9% MSO) higher in the virtual population. Because we knew *a-priori* that 1200 virtual subjects (4.8% of the population) had hotspot intensities more than 10% MSO higher than their ground truth thresholds, a modified fast binary search was also tested, with bounds set to 15% MSO below and 5% MSO above the hotspot intensity. This modification resulted in only six virtual subjects having initial lower bounds above their ground truth thresholds, for which the original fast binary search could never reach threshold.

For all relative-frequency methods, the number of stimuli until the method terminated had a wide distribution (Figs. 1B‒D and S1‒S3, Table S1). Compared to the IFCN five-out-of-ten methods (Fig. 1B, 64 pulses), the staircasing (Fig. 1C, 51 pulses) and binary-search variants (Fig. 1D, default variant 43 pulses and fast variants 31‒ 33 pulses) required fewer pulses on average, in agreement with Boidequin et al. (2026). The binary search variants were also faster on average compared to the staircasing method and had a much tighter distribution of pulse numbers with few outliers. Unlike the IFCN five-out-of-ten methods (*r*^2^ > 0.73), the threshold errors of the variants were also not correlated with the number of pulses required (*r*^2^ < 0.04) (relative errors in Fig. 1B‒D, absolute errors in Fig. S1). However, error distributions of the variants had similar widths as the IFCN method and were also biased albeit with smaller magnitudes (median of relative errors: IFCN 1.07%, staircasing −0.30%, binary search −0.21%, fast binary search 0.48%, and modified fast binary search −0.17%). Using a larger pulse count between ten and twenty pulses at each intensity for the ten-out-of-twenty rule (Rossini et al., 2015; Wang et al., 2023) slightly improved the error distributions for all methods but doubled the number of pulses (Fig. S2‒S3, Table S1).

## 5. Discussion and conclusion

The theoretical analysis showed that relative frequency methods all suffer from the same high probability of false decisions at each intensity as they only include the information of the limited small local sample to estimate a relative frequency in a situation with high variability. Therefore, they cannot guarantee convergence to and accuracy of the threshold. The presented probabilities are still optimistic, as the input–output curve in the analysis (see Eq. 5 in Supplement) has very “light tails” and as such very low probability of unexpected responses, i.e., response at substantially subthreshold intensity or no response at substantially suprathreshold intensity. Actual muscle response distributions are known to have much “thicker tails” (Goetz et al., 2019), which exacerbate the probability of false decisions for relative frequency methods far from threshold. Indeed, the simulation results showed that all relative frequency methods have wide spreads in their distributions of pulse numbers and threshold errors. The variants were able to reduce the average number of pulses required compared to the conventional method, but not the distribution of threshold errors. Indeed, the threshold differences between several relative frequency methods reported by Boidequin et al. (2026) have wide distributions and a significant portion of outliers that can be as large ±10% MSO, which demonstrates their inherent variability and unreliability.

In comparison, more rigorous estimation methods including maximum likelihood estimation (Awiszus, 2003) and stochastic approximation (Wang et al., 2023, 2026b) reach higher threshold accuracy with fewer pulses than relative-frequency methods and their variants. The number of pulses sufficient to obtain accurate thresholds with relative errors of 5% are between twenty to thirty (Figs. 1D_i_ and S1C_i_, and Table S2) and can sometimes be as few as fourteen (Awiszus and Borckardt, 2011; Julkunen, 2019; Wang et al., 2023, 2026b). Therefore, these faster and more accurate methods should become the standard for motor thresholding of TMS, and the use of methods based on relative frequency should be discouraged.

## Acknowledgments

Research reported in this publication is supported by the National Institutes of Health of the United States of America under Award Numbers R01 NS117405, RF1 MH124943, R01 MH129302, and R61 MH129471. The content is solely the responsibility of the authors and does not necessarily represent the official views of the funding agencies.

## Author contribution

BW, SMG, and AVP conceived the study. BW performed theoretical analysis, simulation, data analysis and visualization, and wrote the manuscript. AVP secured funding and resources for the study. All authors revised and approved the final version of the manuscript. Computational support was provided by the Duke Compute Cluster.

## Availability statement

The data that support the findings of this study are openly available at the Duke Research Data Repository with the following URL/DOI (Wang et al., 2026a): https://doi.org/10.7924/r4r551

## Conflict of interest

BW, SMG, and AVP are authors of SAMT, an online software for TMS motor threshold estimation. The copyrights of SAMT are owned by Duke University and the University of Birmingham, and its license for commercial use is managed by Duke University. BW declares no other relevant conflict of interest. SMG has previously received research funding from Magstim as well as royalties from Rogue Research. AVP has received equity options and consulting fees from Ampa; patent royalties and consulting fees from Rogue Research, consulting fees from Magnetic Tides, hardware loans from MagVenture, and hardware donations from Magstim.

## Supplement

### Supplementary methods—false decision probability

For a fixed, even number of pulses, *N*, and at an TMS intensity *x* with an underlying response probability *p*_*r*_(*x*), the probabilities that the relative-frequency method determines the current intensity as subthreshold or suprathreshold after applying *k* pulses respectively are

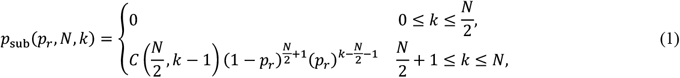

and

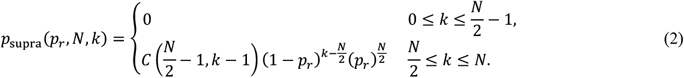

Here, *C*(*N, k*) is the number of *k*-combinations for a set of *N* elements. Hence, the total probabilities that the protocol decides that the current intensity is subthreshold or suprathreshold are given as

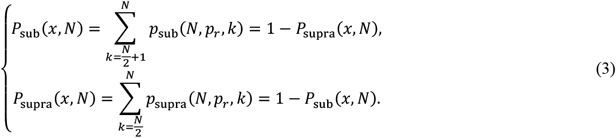

The probability of making a false decision is therefore

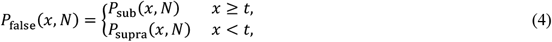

where *t* is the threshold.

For most muscles, previous studies (Awiszus, 2003) describe the response rate *p*_*r*_(*x*) by a sigmoidal function in the form of a cumulative Gaussian distribution

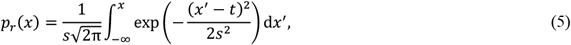

where *s* = 0.07 ⋅ *t* is the average spread parameter that is proportional to the threshold (Fig. 1A, black line). Under these assumptions, the false decision probabilities for some values of *N* are calculated as a function of intensity given relative to threshold (Fig. 1A, colored solid lines).

## Supplementary figures

**Figure S1:**
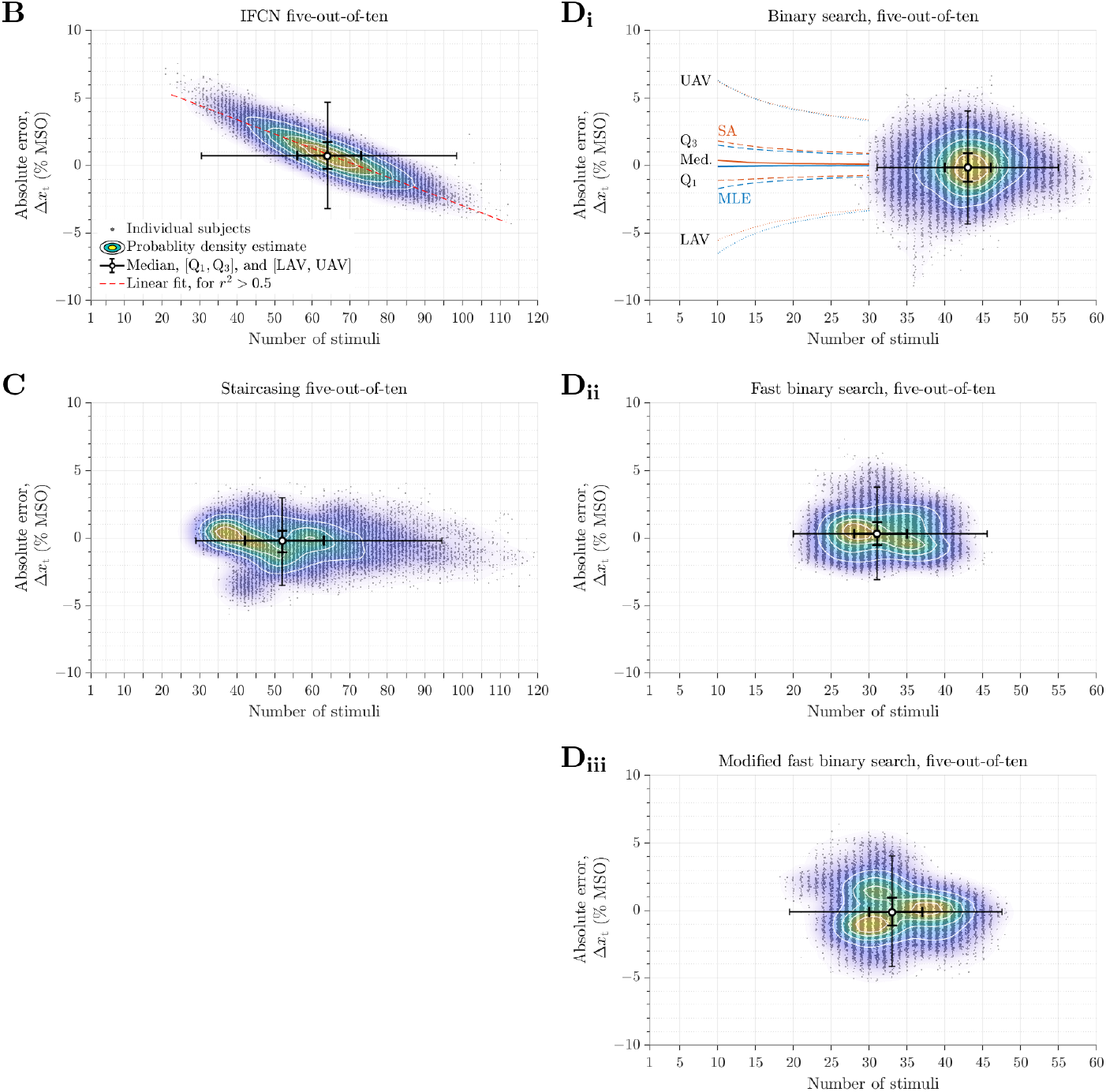
Absolute threshold errors of the relative-frequency methods with five-out-of-ten count. The absolute threshold errors Δ*x*_t_ are shown similarly as the relative threshold errors in Figure 1B–D. Panel A reproduced with data from Fig. S3B_i_ of Wang et al. (2023).

**Figure S2:**
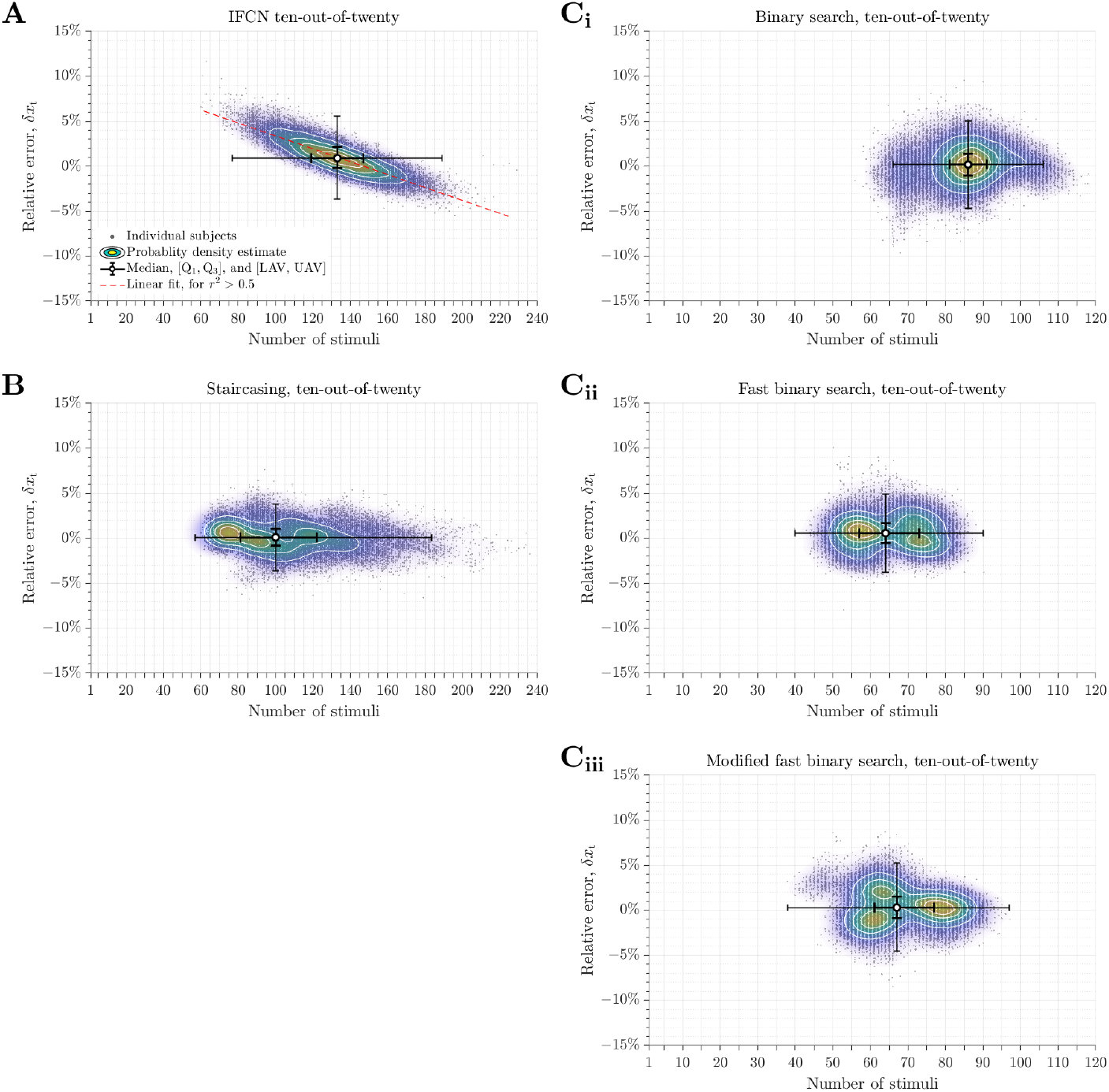
Relative threshold errors of the relative-frequency methods with ten-out-of-twenty count. Shown similarly as the relative threshold errors in Figure 1B–D for five-out-of-ten count. Panel A reproduced with data from Fig. 2A_i_ of Wang et al. (2023).

**Figure S3:**
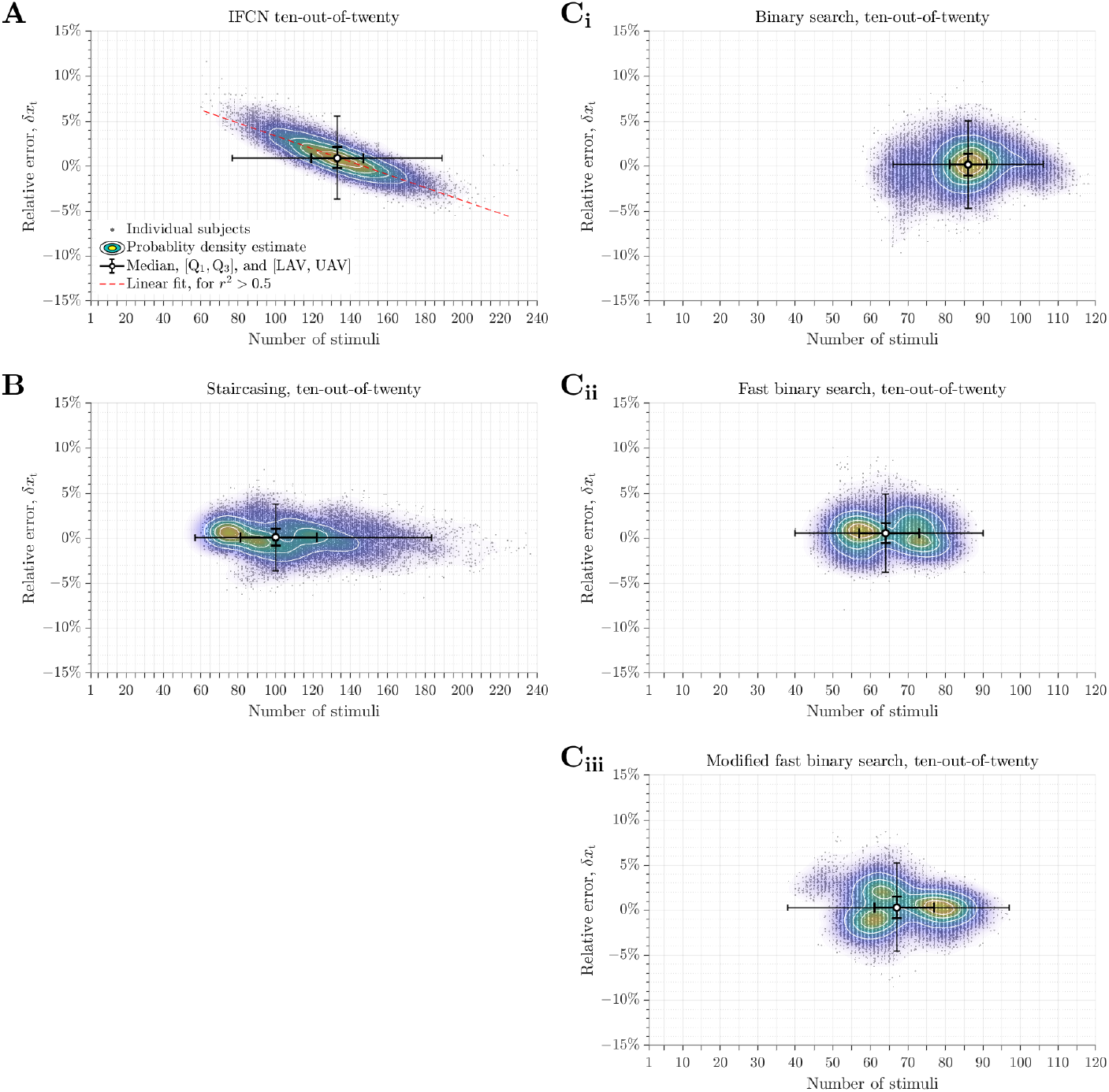
Absolute threshold errors of the relative-frequency methods with ten-out-of-twenty count. Shown similarly as the relative threshold errors in Figure S2. Panel A reproduced with data from Fig. S3A_i_ of Wang et al. (2023).

## Supplementary table

**Table S1.** Statistics of number of stimuli, relative errors, and absolute errors. (related to Figures 1B–D and S1–S3)

| Count | Method | Number of stimuli | | Relative error $\delta x_t$ | | | | $\vert \delta x_t\vert $ | Percentage of $\vert \delta x_t\vert > 5\%$ | Absolute error $\Delta x_t$ (% MSO) | | | | $\vert \Delta x_t\vert $ (% MSO) |
| --- | --- | --- | --- | --- | --- | --- | --- | --- | --- | --- | --- | --- | --- | --- |
| | | Mean ± S.D. | Median | Mean ± S.D. | Median | Slope | $r^2$ | Median | | Mean ± S.D. | Median | Slope | $r^2$ | Median |
| 5/10 | IFCN<sup>#</sup> | 64.5±12.3 | 64 | 1.23%±2.28% | 1.07% | −0.16% | 0.78 | 1.61% | 6.0% | 0.80±1.47 | 0.72 | −0.11 | 0.79 | 1.07 |
| | Staircasing | 54.0±15.1 | 52 | −0.41%±1.94% | −0.30% | −0.02% | 0.02 | 1.22% | 2.0% | −0.27±1.26 | −0.20 | −0.013 | 0.03 | 0.81 |
| | Binary search | 43.0±4.5 | 43 | −0.23%±2.46% | −0.21% | 0.03% | 0.002 | 1.58% | 4.6% | −0.16±1.61 | −0.14 | 0.023 | 0.004 | 1.05 |
| | Fast binary search | 31.7±4.6 | 31 | 0.60%±1.95% | 0.48% | −0.01% | 0.001 | 1.29% | 2.4% | 0.40±1.27 | 0.32 | −0.009 | 0.001 | 0.86 |
| | Modified fast binary search | 33.5±4.9 | 33 | −0.07%±2.40% | −0.17% | −0.04% | 0.006 | 1.56% | 4.2% | −0.04±1.55 | −0.11 | −0.025 | 0.006 | 1.04 |
| 10/20 | IFCN<sup>#</sup> | 133.0±20.7 | 133 | 1.00%±1.75% | 0.91% | −0.07% | 0.73 | 1.28% | 1.8% | 0.66±1.13 | 0.60 | −0.05 | 0.74 | 0.84 |
| | Staircasing | 104.8±27.9 | 100 | 0.08%±1.48% | 0.10% | −0.01% | 0.04 | 0.94% | 0.22% | 0.05±0.96 | 0.06 | −0.007 | 0.04 | 0.62 |
| | Binary search | 85.7±8.9 | 86 | 0.16%±1.89% | 0.18% | 0.03% | 0.01 | 1.21% | 1.4% | 0.09±1.26 | 0.12 | 0.020 | 0.02 | 0.80 |
| | Fast binary search | 64.7±9.0 | 64 | 0.59%±1.63% | 0.56% | −0.004% | 0.0004 | 1.15% | 0.66% | 0.39±1.06 | 0.37 | −0.003 | 0.001 | 0.76 |
| | Modified fast binary search | 68.8±9.7 | 67 | 0.34%±1.84% | 0.28% | −0.01% | 0.004 | 1.23% | 1.0% | 0.22±1.20 | 0.19 | −0.008 | 0.004 | 0.81 |

<sup>#</sup> IFCN methods reproduced with data from Wang et al. (2023)

**Table S2.** Statistics of relative errors, and absolute errors of maximum likelihood estimation and stochastic approximation methods at several pulse numbers. (related to Figures 1D_i_ and S1C_i_), calculated using data from Wang et al. (2023)

| Method | Pulse number | Relative error $\delta x_t$ | | $\vert \delta x_t\vert $ | Percentage of<br>$\vert \delta x_t\vert > 5\%$ | Absolute error $\Delta x_t$ (% MSO) | | $\vert \Delta x_t\vert $ (% MSO) |
| --- | --- | --- | --- | --- | --- | --- | --- | --- |
| | | Mean ± S.D. | Median | Median | | Mean ± S.D. | Median | Median |
| Maximum likelihood estimation | 15 | -0.08%±2.92% | -0.05% | 1.87% | 8.2% | -0.06%±1.89% | -0.04% | 1.24% |
| | 20 | -0.03%±2.44% | 0.00% | 1.57% | 4.1% | -0.02%±1.57% | 0.00% | 1.05% |
| | 25 | -0.02%±2.15% | -0.01% | 1.39% | 2.3% | -0.01%±1.39% | -0.01% | 0.92% |
| | 30 | -0.02%±1.94% | -0.01% | 1.25% | 1.3% | -0.01%±1.25% | -0.01% | 0.83% |
| Stochastic approximation, adaptive stepping | 15 | 0.31%±2.73% | 0.33% | 1.80% | 7.0% | 0.21%±1.77% | 0.23% | 1.20% |
| | 20 | 0.22%±2.33% | 0.23% | 1.54% | 3.5% | 0.15%±1.52% | 0.16% | 1.02% |
| | 25 | 0.18%±2.08% | 0.20% | 1.36% | 1.9% | 0.12%±1.35% | 0.13% | 0.90% |
| | 30 | 0.14%±1.90% | 0.16% | 1.24% | 1.2% | 0.09%±1.23% | 0.11% | 0.82% |

## Notes

https://doi.org/10.7924/r4r551

